# High-speed widefield handheld swept-source OCT angiography with a VCSEL light source

**DOI:** 10.1101/2021.04.10.439295

**Authors:** Shuibin Ni, Xiang Wei, Ringo Ng, Susan Ostmo, Michael F Chiang, David Huang, Yali Jia, J. Peter Campbell, Yifan Jian

**Affiliations:** Casey Eye Institute, Oregon Health & Science University, Portland, OR 97239, USA; Department of Biomedical Engineering, Oregon Health & Science University, Portland, OR 97239, USA; Department of Engineering Science, Simon Fraser University, Burnaby, Canada; National Eye Institute, National Institutes of Health, Bethesda, MD 20892, USA

## Abstract

Optical coherence tomography (OCT) and OCT angiography (OCTA) enable noninvasive structural and angiographic imaging of the eye. Portable handheld OCT/OCTA systems are required for imaging patients in the supine position. Examples include infants in the neonatal intensive care unit (NICU) and operating room (OR). The speed of image acquisition plays a pivotal role in acquiring high quality OCT/OCTA images, particularly with the handheld system, since both the operator hand tremor and subject motion can cause significant motion artifacts. In addition, having a large field of view and the ability of real-time data visualization are critical elements in rapid disease screening, reducing imaging time, and detecting peripheral retinal pathologies. The arrangement of optical components is less flexible in the handheld system due to the limitation of size and weight. In this paper, we introduce a 400-kHz, 55-degree field of view handheld OCT/OCTA system that has overcome many technical challenges as a portable OCT system as well as a high-speed OCTA system. We demonstrate imaging premature infants with retinopathy of prematurity (ROP) in the NICU, and patients with incontinentia pigmenti (IP) in the OR using our handheld OCT system. Our design may have potential for improving the diagnosis of retinal diseases and help provide a practical guideline for designing a flexible and portable OCT system.

## 1. Introduction

Optical coherence tomography (OCT) and OCT angiography (OCTA) have become an essential tool for health evaluation of the retina in adults and are being explored as a noninvasive method that can produce objective biomarkers, which leads to earlier and more objective diagnosis in adult retinal diseases. OCT has also become part of the standard of care in adult retinal practices due to the value added by the ability to identify subclinical disease features and objectively track disease progress in many of the leading causes of blindness [1–4]. Treatment paradigms for diabetic retinopathy and age-related macular degeneration now integrate OCT findings into the disease management. OCTA is based on detecting flow-related amplitude and/or phase variation/decorrelation in successive B-frames acquired at the same location. The intrinsic motion contrast obviates the needs for the injection of any extrinsic contrast agent. OCTA research and clinical applications are expanding greatly due to the ability to noninvasively visualize the retinal and choroidal circulation in adults [5–8]. Although OCTA has been demonstrated since 2006 [9], clinical applications require many technical improvements to enhance practicality. Most of the OCTA methods require at least two repeated OCT B-scan at the same transverse location to detect blood flow, which significantly increases the image acquisition time and motion artifacts. Also, OCTA is more sensitive to transverse digital resolution, optical resolution, a large imaging spot size, and low sampling rate may result in OCTA images with poor contrast and signal-to-noise ratio (SNR). In the commercial OCT/OCTA systems, the OCT A-scan rate is typically lower than 100-kHz, thus a compromise has to be made between acquisition time and field of view. Most commercial OCT/OCTA imaging devices are desktop systems that are limited in speed and designed for adults with their ability to fixate. The benefits of OCTA have yet to extend to general pediatric retina as it is difficult and impractical to image unanesthetized and uncooperative infants with a desktop OCTA system. Even the commercially available portable devices for young children are unsuitable for OCTA due to insufficient speed.

Several groups, including our own, have developed prototype handheld OCT systems that can be used to image patients in the supine position [10–13]. And the improved portability allows screening to be done within primary care settings or the operating room. The development of swept-source OCT sparked rapid increase of OCT imaging speed from several tens of kilohertz (kHz) to more than one megahertz (MHz), making OCTA with handheld probe feasible. Campbell *et al.*, Viehland *et al.* and Song *et al.* demonstrated handheld OCTA for imaging neonates with retinopathy of prematurity (ROP) [11,12,14], a retinal vasculature disease that affects premature babies and could lead to permanent vision loss if not diagnosed and treated in time. Several clinical studies showed the potential of using OCT and OCTA to study ROP [15–18]. Nadiarnykh *et al.* showed clinical utility of a handheld OCTA system for diagnostic imaging of pediatric retinoblastoma as well as monitoring disease progression and response to treatment [19]. These prototype handheld systems did not have the field of view that is needed for visualizing peripheral retina where the pathologies in diseases such as ROP occur. We are the only group to have demonstrated the ability to obtain ultra-wide field (UWF) OCT images on non-sedated babies with ROP [14]. Due to the challenges of imaging a non-fixating and non-sedated neonate, it has been difficult to consistently obtain high quality data for biomarker evaluation in ROP. In addition to the hardware constraints, current research and commercial handheld OCT systems lack the ability to visualize the retina in real time due to the intensive computational requirement for OCT image processing. In comparison, for the desktop OCT systems for imaging adults, a simple preview of selected OCT B-scans is often sufficient to evaluate and optimize image quality. Patient alignment and locating landmarks on ocular fundus can be easily accomplished with the help of fixation target and a secondary fast fundus imaging modality such as a scanning laser ophthalmoscope or a near infrared fundus camera. In handheld OCT, pupil camera and on-probe display have been integrated to facilitate alignment [12,20,21]. However, these features may add complexity and weight in the handheld system. Real-time visualization of the OCT *en face* and cross-sectional scans on the imaging probe could be instrumental for guiding the imaging procedure, confirming the areas of interests captured and reducing imaging time for the most fragile patient populations.

There is a need for a dedicated portable handheld OCT/OCTA retina scanner for the pediatric population that extends the field of view to the peripheral retina and displays the cross-sectional and *en face* views of the targeted area in real time. In this manuscript, we demonstrate a 400-kHz, 55-degree field of view handheld OCT/OCTA system that has overcome many technical challenges in high-speed OCTA imaging as well as in pediatric retinal imaging in general and shows its potential to improve diagnosis of retinal diseases and provide clinical validation.

## 2. Methods

### 2.1 OCT system design

The schematic of the high-speed and widefield handheld OCTA system is shown in Fig. 1. The swept-source laser used in our system was a vertical cavity surface emitting laser (VCSEL) (SVM10F-0210, Thorlabs, Inc., USA). This laser has an A-scan sweep rate of 400-kHz, a center wavelength of 1060 nm and 100 nm bandwidth, corresponding to a theoretical axial resolution of 4.94 μm in the air. The high output power of this VCSEL source allows us to optimize the fiber-based interferometer so that we could deliver sufficient and safe amount of imaging beam power to the sample arm while maximizing collection efficiency. The 4-coupler interferometer in this system consisted of a 10/90 fiber coupler, a 20/80 coupler, and two 50/50 fiber couplers. The OCT sample arm was connected to a set of 10/90 coupler and a set of 20/80 coupler. The 10/90 coupler split 90% of the laser output power to the 20/80 coupler, and the 20/80 coupler split 20% of its input power to the sample arm. The 10% laser output power that was split by the 10/90 fiber coupler was delivered into the reference arm through a 50/50 fiber coupler. 80% of the light collected from the sample arm and the 50% of the light reflected from reference arm were combined within the 50/50 fiber coupler and detected with a balanced detector (PDB482C-AC, Thorlabs, Inc. USA).

**Fig. 1.**
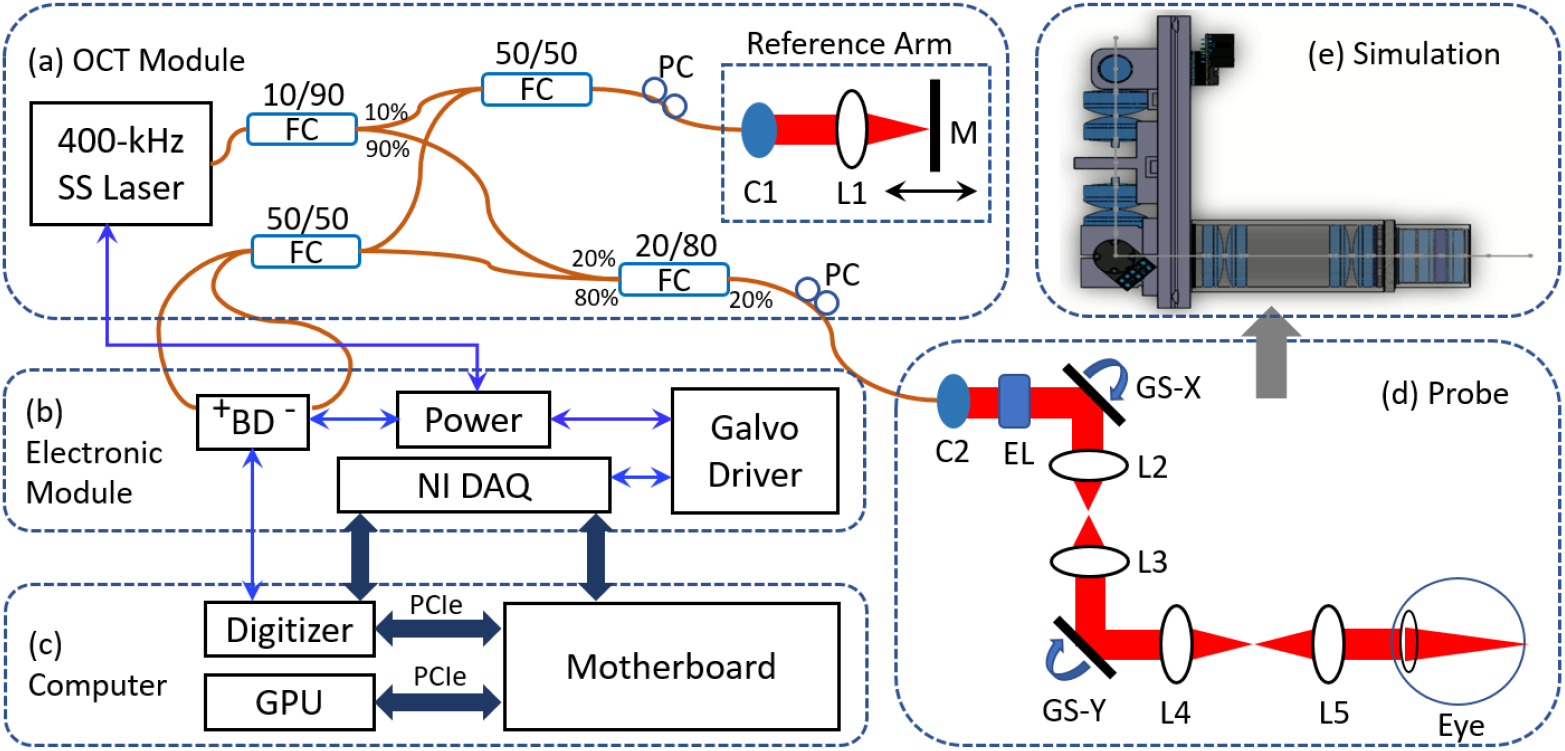
Schematic diagram of the handheld OCTA system. (a) OCT module, including the laser source, the spectrometer, and the motorized reference arm; (b) Electronic module, including the VCSEL source, the balanced detector, data acquisition, control module (BNC-2110, National Instrument, Inc. USA) and digital servo driver for galvanometer scanner (Pangolin Laser System, Inc. USA); (c) Custom-built computer integrated with GPU and digitizer (ATS9373, Alazar Technologies, Inc. Canada) via PCI Express (PCIe) link; (d) Optical layout of the probe; (e) Opto-mechanical simulation of the probe in SolidWorks. M: mirror; PC: polarization controller; FC: fiber coupler; BD: balanced detector; GPU: graphics processing unit. GS-X: the fast axis of galvanometer scanner; GS-Y: the slow axis of galvanometer scanner; L1-L5: lens; C1, C2: collimator; EL: electronically focus tunable lens.

A 3 m fiber patch cord was connected to the sample arm for flexible handheld operation and a 3.3 m fiber patch cord was connected to the reference arm to match the optical path length and dispersion of the sample arm. In addition to matching the optical path length, the polarization state between the sample and reference arm needed to be matched for interferometry. Bending of the optical fiber patch cord in the sample arm would alter the polarization state, and the polarization mismatch between the sample and reference arm would decrease the OCT signal to noise ratio significantly. A two-paddle manual fiber polarization controller with the loop diameter of 18 mm (FPC020, Thorlabs, Inc., USA) and a two-paddle motorized fiber polarization controller (MPC220, Thorlabs, Inc., USA) were installed in the path of sample arm and reference arm, respectively. A custom-built motorized reference arm with tunable range of 100 mm was implemented to adjust the optical delay when switching between adults and neonates scanning.

### 2.2 Optical design of the handheld OCTA probe

The optical design of the handheld OCTA probe is shown in Fig. 2. The sample arm optics were designed and optimized in OpticStudio (Zemax, LLC, USA), and assembled with off-the-shelf optical components. The distances between the optical elements in the sample arm were optimized by minimizing the spot radius on the focal plane and the imaging beam wander on the pupil plane. Starting from the 20/80 fiber coupler, the beam entered through a 3 m single mode custom patch cable and was collimated into a 2 mm beam by a reflective collimator (RC02APC-P01, Thorlabs, Inc., USA). It then passed through an electronically focus tunable lens (EL-3-10, Optotune, Switzerland), a pair of separated galvanometer driven mirrors (Saturn 1B, ScannerMax, USA) with a set of achromatic lenses set up in the 4f configuration in between to relay the beam. The scanning beam after the slow axis of galvanometer scanner (GS-Y, Fig. 2b) entered a pair of achromatic doublet lenses with 30 mm diameter and 100 mm focal length as scan lens (L5-L6, Fig. 2b), and then entered a 5-lens group that together to form an ocular lens (L7-L11, Fig. 2b). The 5-lens group consisted of a pair of 30 mm positive achromatic doublet lenses with 100 mm focal length, 1 inch negative achromatic doublets lens with −100 mm focal length, and a pair of 1 inch positive achromatic doublet lenses with 50 mm focal length. The scan lens and ocular lens worked as 1.6× telescope, which amplified the scanning angle from 34° to 55°. The imaging beam was also demagnified from 2.1 mm in the slow axis of the galvanometer scanner to 1.3 mm at the pupil plane. The working distance of the handheld probe was 9.8 mm, which was reasonable for imaging both the adults and neonates.

**Fig. 2.**
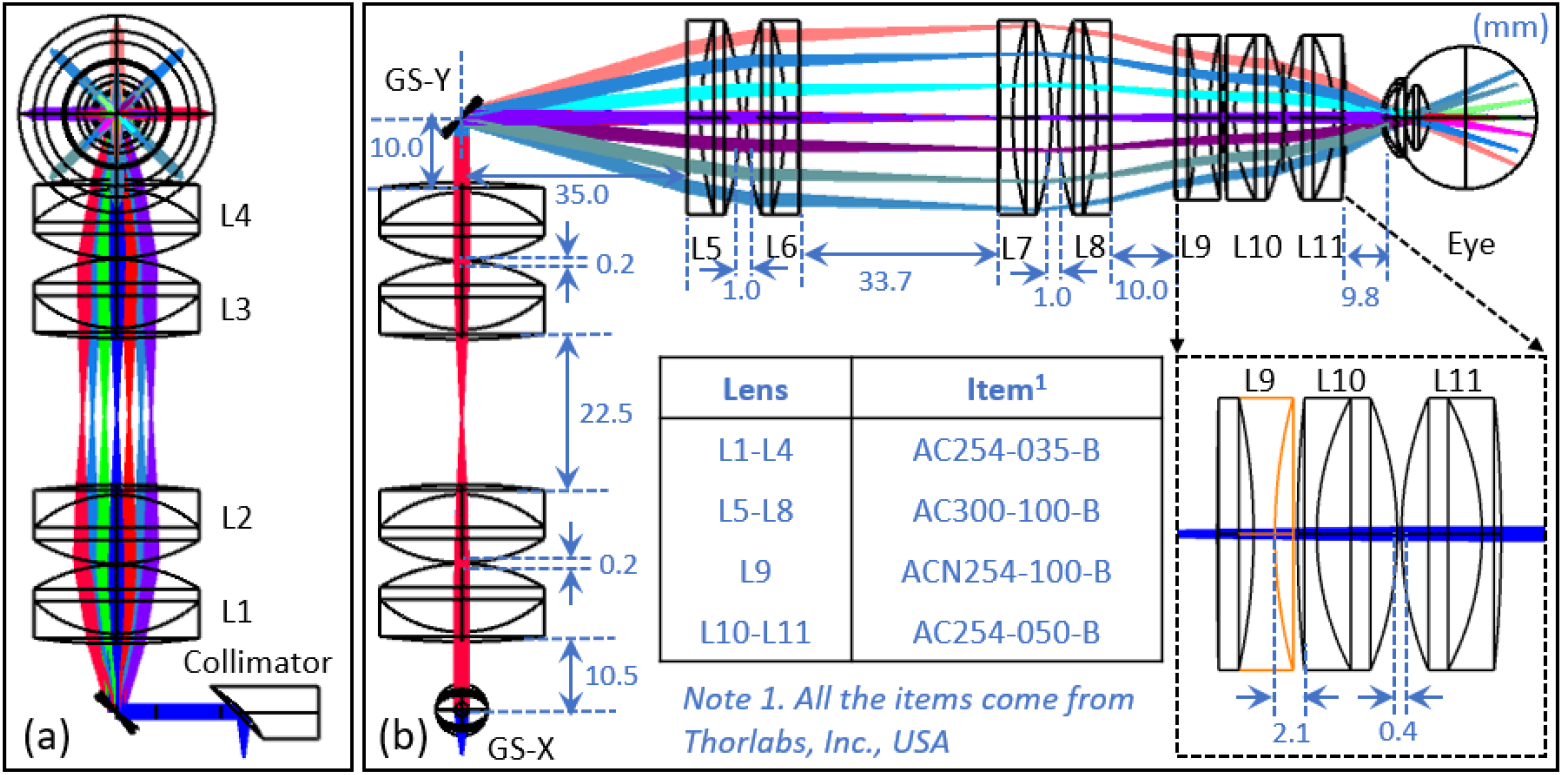
(a) Side view of 3D layout in OpticStudio. (b) Front view of 3D layout in OpticStudio. All the lenses configurations in the probe and the distance between components were illustrated.

The fast and slow axis of the galvanometer scanning mirrors were separated to reduce the vignetting artifacts and the imaging beam wander on the pupil plane a technique often used in adaptive optics retinal imaging system [22,23]. The ocular lens was made up of a group of 5 lenses with different diameters to increase the scanning angle and improve the optical performance. For comparison, we simulated a second optical design, which was referred to as the ‘conventional handheld OCT system’ (Fig. 3b and Fig. 3d). In this classical optical design of retinal OCT system, the galvanometer scanning mirrors were not separated and the ocular lens (5-lens group) was replaced with a pair of achromatic doublet lenses with a 30 mm effective focal length to ensure the same beam size at the pupil plane compared to our design. Fig. 3a and Fig. 3b show the comparison of multi-configuration spot diagram in these two optical designs with the same field of view. These figures demonstrated that the aberration at large scanning angles was significantly reduced after implementing the design improvement. The root mean square (RMS) of the spot radius of the upper left corner (the purple ray) in Fig. 3a and Fig. 3b was decreased from 21.527 μm to 15.821 μm after optimization. In addition to the focused spot radius on the imaging plane which represents the transverse resolution, the imaging beam wander on the pupil plane is also a critical and yet often omitted metric in evaluating the optical performance of a retinal OCT imaging system. It is especially important in widefield OCT system as the vignetting and aberration are introduced significantly at larger scanning angles [24]. Fig. 3c and Fig. 3d show the comparison of imaging beam wander on the pupil plane in these two optical designs with 0°, 20° and 38° scanning angles. The 20° angle translates to a 6×6 mm field of view in an adult normal eye. The 38° is the maximum angle in the conventional handheld OCT system. The results of the comparison indicated that the integrated galvanometer scanning mirrors layout had a great impact on the beam footprint on the pupil plane. The maximum diameter of the footprint diagram in the optimized handheld OCT system and the conventional handheld OCT system was 1.8788 mm and 2.4122 mm, respectively. The amount of imaging beam wander on the pupil plane was reduced significantly after separating the galvanometer scanning mirrors because both galvanometer mirrors are optically conjugated on the pupil plane. To further increase the scanning angle, 30 mm diameter rather were selected over typical 1 inch lenses for L5-L8. The larger diameter of the lenses closes to the scanner with the same effective focal length, the wider scanning angle was achieved. One trade-off for using larger lenses was that it increased the weight of the probe, which may cause operator fatigue. Another disadvantage of implementing larger lenses was that it disrupted the ergonomics of the handheld probe, making it more difficult to hold steady and maneuver. A shorter effective focal length at this position could also increase scanning angle, but the imaging beam size on the pupil plane would be further demagnified, which reduced the system’s numerical aperture and transverse optical resolution. This issue could be solved by choosing a collimator with longer focal length; however, corresponding larger aperture galvanometer mirrors are needed to accommodate the increased beam diameter. Because this system’s B-scan frame rate is in the order of kHz, it demands high performance from the galvanometer scanning mirror, and in general smaller aperture galvanometers are faster.

**Fig. 3.**
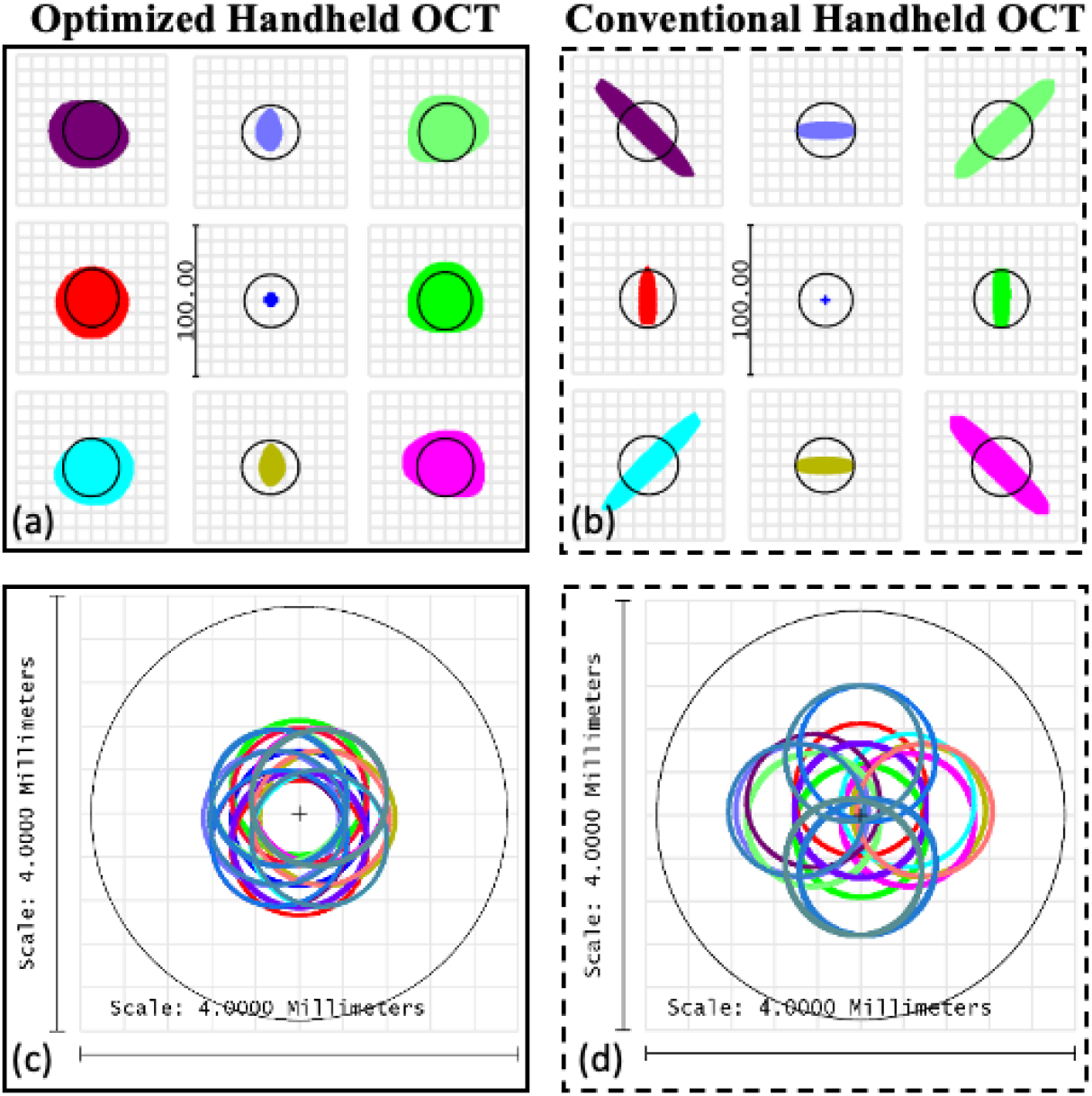
Comparison of multi-configuration spot diagrams for the optimized optical design (a) and conventional optical design (b) of handheld OCT system with a 28° field of view, which is equivalent to 9×9 mm. The radius of Airy Disk (black circle) is 19.02 μm (a) and 19.64 μm (b). Comparison of imaging beam wander on the pupil plane with the combination of 0°, 20° (6×6 mm), and 38° (maximum scanning angle of the conventional handheld OCT system) field of view for the optimized optical design (c) and conventional optical design (d) of handheld OCT system. The maximum radius of the footprint diagram is 0.9394 mm (c) and 1.2061 mm (d).

The primary application of our handheld OCT/OCTA system is screening retinal disease for infants whose eyes are significantly shorter than an adults’ and there is a large variance between premature babies and babies that were born later in the pregnancy terms. Our sample arm’s optical design aims for a balance between the field of view, transverse optical resolution, and performance of the galvanometers. The final beam size on the pupil plane was 1.3 mm, and when imaging an infant with an axial eye length of 17 mm, it results in a focused spot of approximately 12.6 μm in diameter (1/e^2) on the retinal plane.

### 2.3 Mechanical design of the handheld OCTA probe

The optical simulation of sample arm optics was exported to CAD software (Dassault Systèmes, France) for the mechanical design, where the precise locations of each optical components were specified according to the optical simulation. Since the measurement of OCTA was based on the change of blood flow signal, the subtle motion of the system or the subject could affect the results. Compared to the conventional desktop-based OCT system, the motion of the operator’s hand is an extra source of motion artifacts that could corrupt the OCTA signals in the handheld system. Our mechanical design of the handheld imaging probe is ergonomic. With a single hand, the operator could hold the lens tube instead of the body of the probe, which allowed the operator to rest parts of their palm gently on the infant’s forehead and manipulate the eye position using a depressor with their other hand when necessary. The grip permitted the operator to align the target area with a slight rotation of the wrist. The mechanical design for the probe was shown in the Fig. 4.

**Fig. 4.**
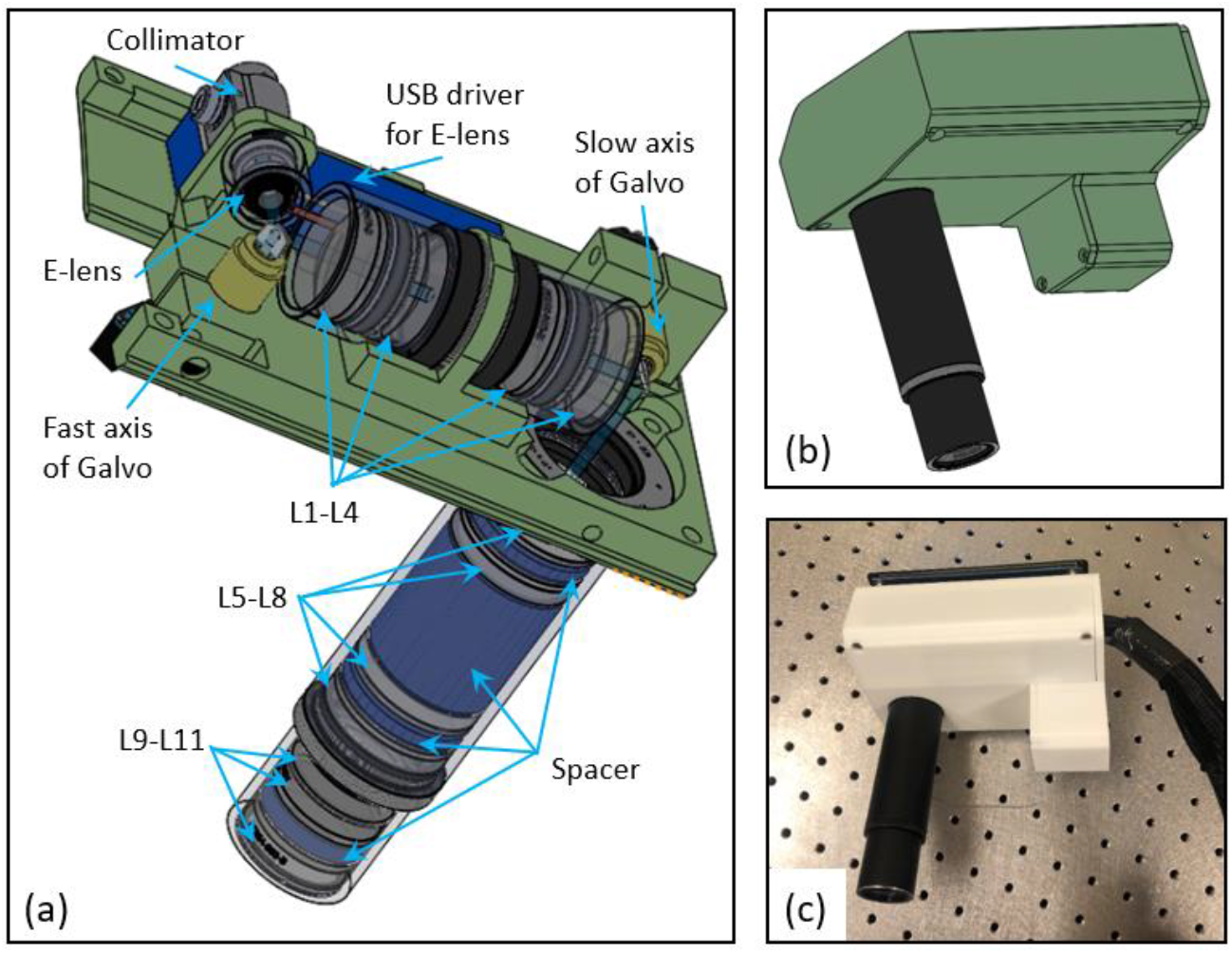
(a) Mechanical layout of the key components of the probe. (b) Appearance of the fully assembled probe in SolidWorks. (c) Photograph of the probe with 3D printed enclosure.

To ensure precise alignment of the lenses, the lens tube with required length was customized from the manufacturer (Thorlabs, Inc., USA). The spacers between the lenses were designed and fabricated according to the optical simulation. Multiple positioning slots were designed on the enclosure to make sure that the components were assembled in the specified location. The enclosures and spacers were fabricated by a 3D printer (S5, Ultimaker B.V., Netherlands) with resolution of ±0.1 mm. The collimator and galvanometer scanners were directly mounted onto the high-resolution 3D printed enclosure without the aluminum heatsink to reduce the bulk and weight of the probe. The last group of lenses required frequent cleaning; therefore, an adapter was used to connect two lens tubes so that it could easily be if necessary. Fig. 4c shows a photograph of the fully assembled handheld probe. The handheld probe, excluding the fiber and electrical cables, weighed about 700 g and with the dimensions of 148 × 97 × 175 mm (length × width × height). A 5.5 inch AMOLED display was integrated on top of the probe to facilitate alignment. Two separate imaging windows, B-scan and *en face* OCT, were shown in the screen (Fig. 5c). All the electrical cables and optical fiber were wrapped into nylon protective sleeve sheath, which provided some minor strain relief on the cables and protected them from abrasion.

**Fig. 5.**
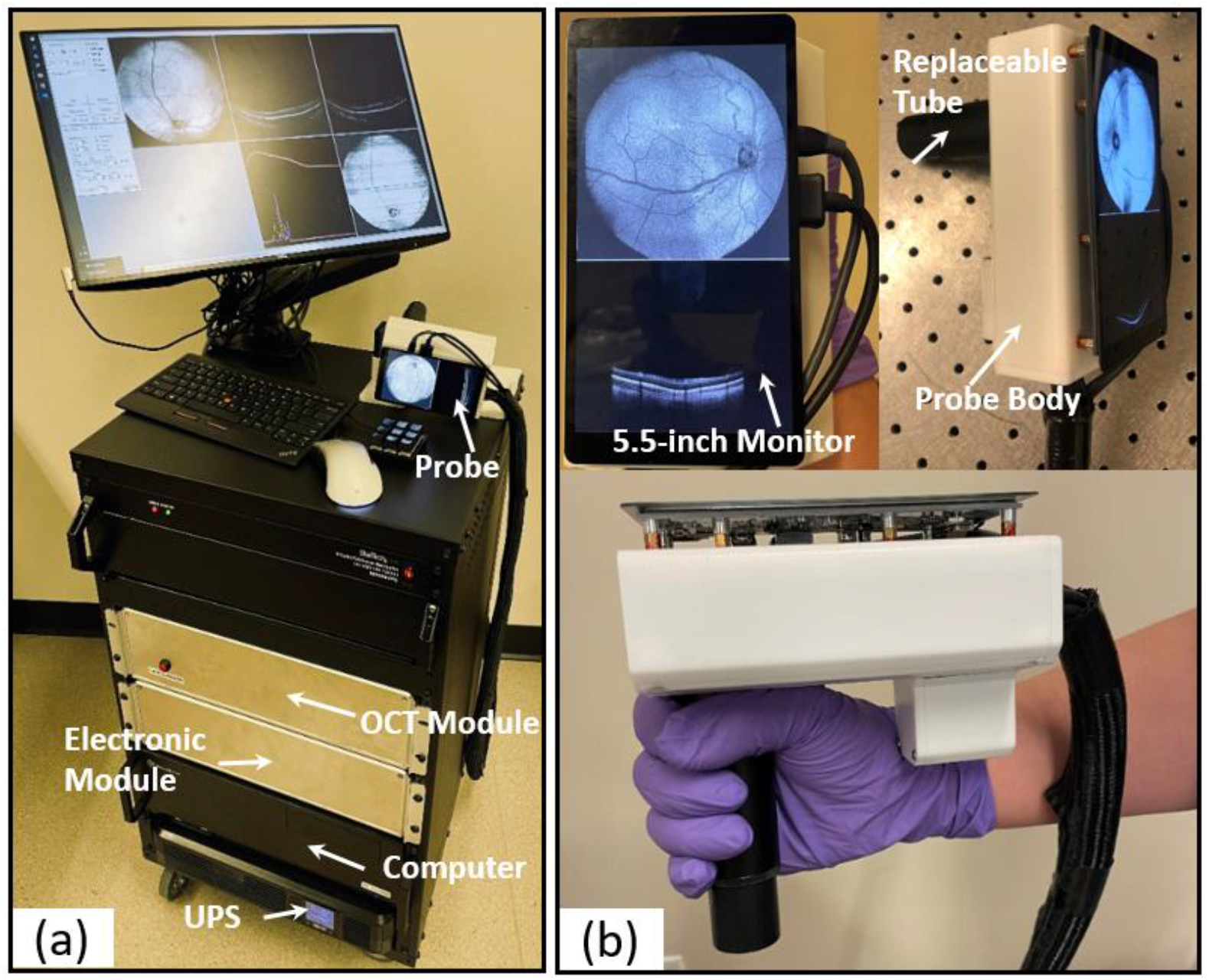
(a) Photograph of the front of fully assembled handheld OCTA system in portable cart. (b) Photograph of the handheld probe.

### 2.4 System assembly

The imaging system consisted of an OCT module, an electronic module, a computer and the handheld probe. The OCT module and electronic module were separately assembled into two 19 inch rack enclosures of 3 rack units (U) in height (EC3U, Thorlabs, Inc., USA). The computer was assembled in a 19 inch rack enclosure of 4U height. The three enclosures were communicated through detachable cables and installed in a 16U server rack with lockable caster wheels (Fig. 5a), which could be transported easily between different sites for imaging. The fully assembled system is shown in Fig. 5. The uninterruptible power supply (UPS), with line-interactive topology, provided backup power to keep the system running during transportation from one ward to another and protected the system from the power fluctuations. The smart cooling fan mounted on the back of the portable cart was taken feedback from a temperature sensor placed near the VCSEL (laser) module to maintain its temperature below 50 °C. To help maintain the proper airflow, panel spacers with perforated venting were installed on the back of the cart.

### 2.5 Image acquisition protocol

The output of the balanced detector was digitized by a 12 bit waveform digitizer board with an external k-clock provided by the laser. The VCSEL source has a long coherence length over 100 mm, so the limiting factor for the OCT imaging depth in our system is the interferogram sampling density. To increase the imaging depth, which is critical in ultrawide field imaging, we used the dual edge sampling feature of the digitizer. This feature allowed us to sample the spectrum at both rising and falling edges of the sampling clock (external k-clock). The 400-kHz VCSEL swept-source laser used in our system has a 50% duty cycle and each raw A-scan spectrum was sampled with 2048 points corresponding to a peak sampling frequency of 1720 MHz and a maximum imaging depth of 6 mm in air. The total round trip optical path length in our system is 12.97 m in air, which created a 43.22 ns delay between the external k-clock signal and the interferometric signal when it was sampled at the digitizer. To counteract this delay, a 7.62 m (25 feet) coaxial cable was added between the output of the k-clock signal on the laser module and the input external clock port on the digitizer. The remaining mismatch was fine-tuned by the laser’s software.

High-speed swept-source OCT significantly reduces motion artifacts for *in vivo* clinical imaging. It is especially critical to handheld clinical imaging applications. OCTA is based on detecting flow-related amplitude and/or phase variation/decorrelation in successive B-frames acquired at the same location. The intrinsic motion contrast obviates the need for the injection of any extrinsic contrast agent; however, a 3-5 ms interval [25] between the repeated B-scans is desired for OCTA to detect sufficient blood flow coming from motion contrast. In high-speed OCT, the interval between consecutive B-frames at the same position is too short to detect moving red blood cells. Furthermore, the B-frame rate is limited by the frequency of the scanner (<500 Hz at large scanning angles). Instead of slowing the system to achieve the desired 3-5 ms interval [25] to detect capillary blood flow with OCTA, we recently developed a stepped bidirectional scanning pattern [26,27] that enables faster B-frame rate (up to 1 kHz at 30°). In this project, we designed an OCTA scanning pattern (Fig. 6) that performs three repeated B-scans at the same location with 500 A-scans per B-scan and 5 ms interval between the repeated B-scans, leading to a total of 1500 B-scans per volume. Each volume acquisition time was 1.875 s. OCTA B-scans were generated by the split-spectrum amplitude-decorrelation algorithm (SSADA) [28,29], computed from the three repeated B-scans acquired at each position. *En face* OCTA images were generated after manual retinal layer segmentation. The *en face* OCTA image was then processed using medium subtraction method to remove small motion artifacts, a Gabor filter was also applied to enhance the OCTA signal [30,31]. A customized color map was applied for each *en face* image to improve the OCTA visibility. In SSADA, the speckle decorrelation of OCT B-scans generated for different spectral bands of interferometric signal yielding the OCT B-scan images are averaged, improving the signal-to-noise ratio of flow detection over full-bandwidth speckle decorrelation measurement.

**Fig. 6.**
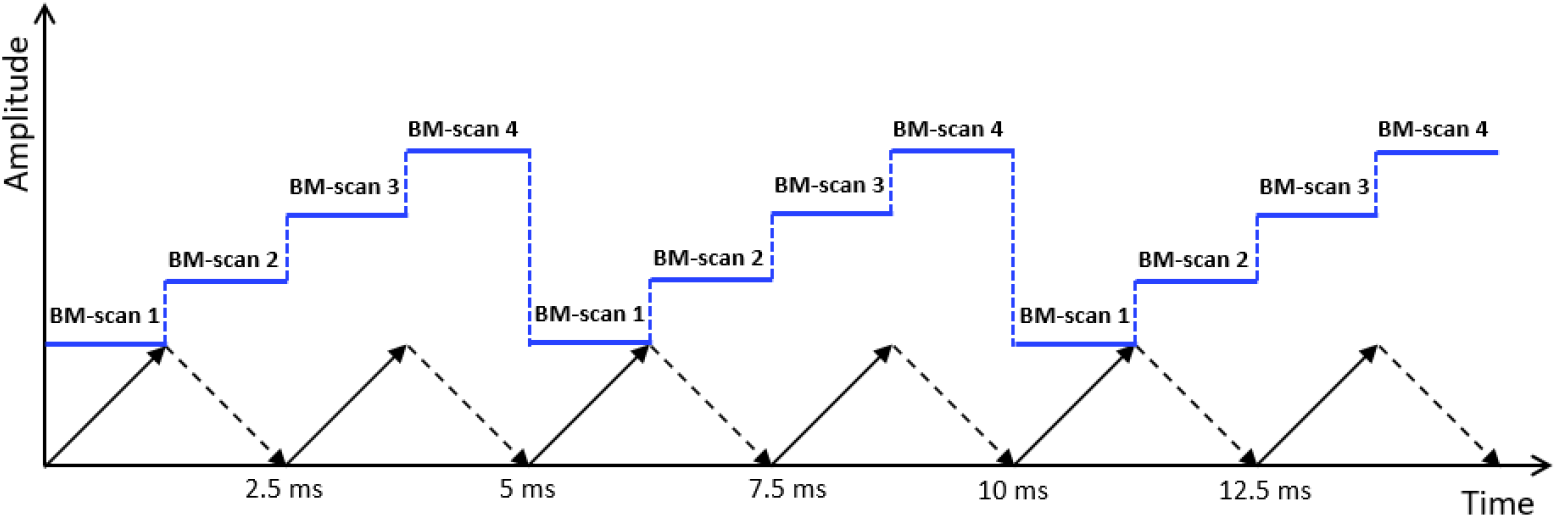
Stepped bidirectional scanning pattern with three repeated B-scans. Solid and dashed arrows represent fast scan (B-scan). Here, three repeated B-scans were used to detect OCTA. The interval between the repeated B-scans was 5 ms.

### 2.6 Software operation and data visualization

OCT and OCTA images were acquired and processed in real-time continuously by our custom written GPU accelerated software OCTViewer, providing an immediate evaluation of the image quality [32,33]. The data acquisition computer consisted of an Intel Core i9 9900k CPU, a NVIDIA Titan RTX GPU, a multifunction I/O DAQ (PCIe 6353, National Instrument, Inc., USA) and a high-speed digitizer (ATS9373, Alazar Technologies, Inc., Canada). In addition to the 5.5 inch display installed on the handheld probe, a 24 inch monitor was also mounted on the imaging cart to display more information, including OCTA *en face*, cross-sectional views, and selected raw and processed A-scans. There were two scanning modes available on the handheld OCTA system. One mode was the warp scan, which was a high-speed alignment mode and used to identify and align the target area. In this mode, *en face* view of the aiming area was visualized continuously. The acquisition time of each volume was 120 ms with 400 A-scan per B-scan and 120 B-scans per volume, which provided real-time feedback in optimizing images. Once the target area on the ocular fundus was located, the focus could be optimized automatically based on the brightness of the *en face* images with a hill-climbing algorithm within 1 second [34]. The second scanning mode was the OCTA mode, which was used to acquire high resolution OCTA volume. A secondary keyboard was connected to the computer with 6 shortcut keys programmed to save data, toggle scanning modes, auto-focus, auto-polarization control, and adjust reference arm length.

### 2.7 Study subjects

Handheld OCT/OCTA retinal imaging was performed on adults, sedated infants, and fully awake neonates. These patients were recruited from the Casey Eye Institute at the Oregon Health & Science University (OHSU). Written informed consent from the patient or parent/guardian in case of minor patients was obtained prior to initiating the study. The research was approved by the Institutional Review Board/Ethics Committee of OHSU in accordance with the Declaration of Helsinki. The imaging for the non-sedated infants was conducted with pharmacological dilation per standard protocols in the operating room and neonatal intensive care unit. The optical power incident on the subject cornea was set to less than 1.8 mW, which is within the American National Standards Institute (ANSI) Z136.1-2000 Standards for the safe ocular laser exposure limit at 1060 nm [35].

## 3. Results

In this study, 3 healthy adult volunteers, 16 awake neonates in the neonatal intensive care unit and 7 pediatric patients under general anesthesia prior to retinal surgery in the operating room were imaged. Prior to the imaging session, the subjects were dilated with cyclomydril, an eyelid speculum was used to keep the eye open and balanced salt solution was applied regularly to prevent the eye from drying. The operator began by sitting close to the head of the neonate and positioned the device close to the eye with the lens cap attached then initiated the automated polarization state matching based on the brightness of the lens cap *en face* image. Next, the imaging session began, starting in warp scan mode, which offers *en face* view at a high refresh rate allowing the operator to quickly align the system to the retina. The reference arm length was predetermined based on the age of the subject and fine-tuned during the imaging procedure. The warp scan mode provided the *en face* view from the front of eye into the retina as the operator adjusted the working distance. Ocular funds would start to appear in the e*n face* view as the probe got closer to the eye, and the operator can identify the larger vascular structures and any extraretinal pathology within approximately 100 × 100 window by pivoting the probe. Once the desired region of interests was located, autofocusing was performed in 1 second based on the brightness of the *en face* view. We then switched to OCTA mode that continuously acquired volumes every 1.875 seconds. The OCT *en face* and B-scan was viewed in the on-probe display and desired volumes were saved in an independent thread so that the image acquisition and processing did not halt during the saving process. Fig. 7 indicates a screen capture of external monitor on the imaging cart demonstrating the data acquisition process described above (Visualization 1).

**Fig. 7.**
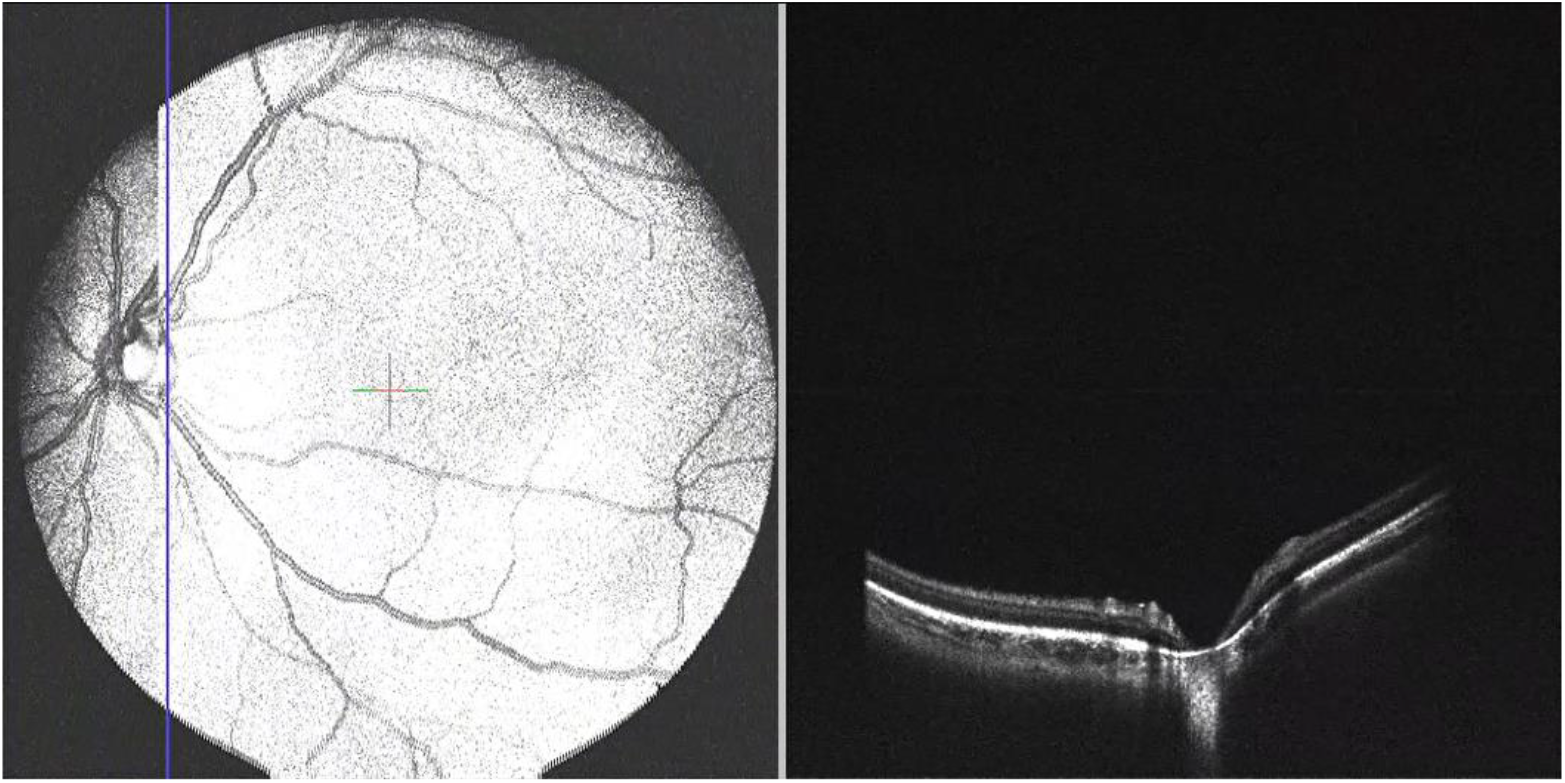
Screen capture of the OCTViewer software representing the data acquisition process.

The retinal OCT/OCTA images from three representative cases-a patient with ROP, a patient with X-linked retinoschisis (XLRS), and a patient with incontinentia pigmenti (IP)-are shown in Fig. 8-11 separately. Fig. 8 shows the *en face* OCT image and corresponding OCTA image of the inner retina from a premature infant with ROP after laser treatment. While laser treatment is often effective in inducing regression of extraretinal neovascularization, in some cases vitreoretinal traction can worsen leading to vascular dragging, retinoschisis, and retinal detachment [36]. These changes are much easier to appreciate on OCT compared to the clinical exam, and early detachment of the retina is typically located in the retinal periphery, out of the field of view of traditional OCT even in adults. Therefore, an imaging system with wide field of view and can visualize the peripheral retina is needed, to follow up after the treatment. Fig. 8a indicates the *en face* OCT image in the peripheral retina. The area indicated by the red arrow (Fig. 8a) is the chorioretinal scar left by the laser treatment, without evidence of residual neovascularization, fibrosis or traction.

**Fig. 8.**
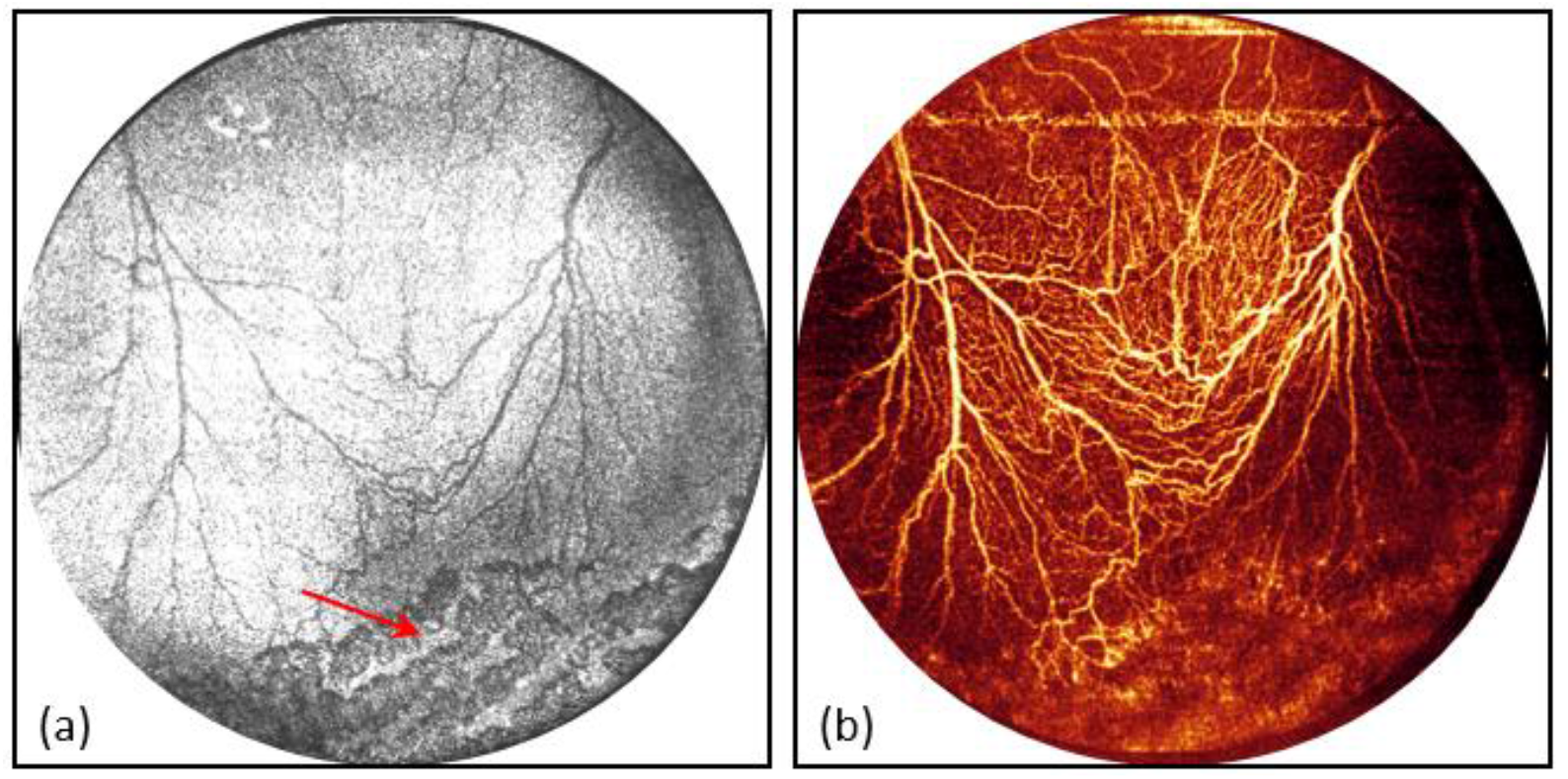
(a) *En face* OCT image from an infant with retinopathy of prematurity (ROP) after laser therapy. Laser scars are shown with the red arrow. (b) Corresponding OCTA *en face* image of the inner retina.

Fig. 9 shows *en face* OCT image, corresponding OCTA *en face* image of the inner retina and two selected B-scans from a patient with XLRS imaged in the OR prior to surgery. The locations of selected B-scans in Fig. 9c are indicated by the blue arrow (①) and green arrow (②) in *en face* OCT image (Fig. 9a). XLRS is a kind of congenital retinal diseases, which caused by genetic mutation on the X chromosome that prevents encoding the protein necessary for the cell-to-cell adhesion throughout the retinal development [37]. Two selected B-scans demonstrate the typical findings of intraretinal schisis, and vitreoretinal traction from a large schisis cavity. One of the key management decisions is based on the distinction between retinoschisis and retinal detachment. Our present system could be used to examine the suspected area and visualize the pathology in real time and higher resolution than is available with other modalities such as ultrasound.

**Fig. 9.**
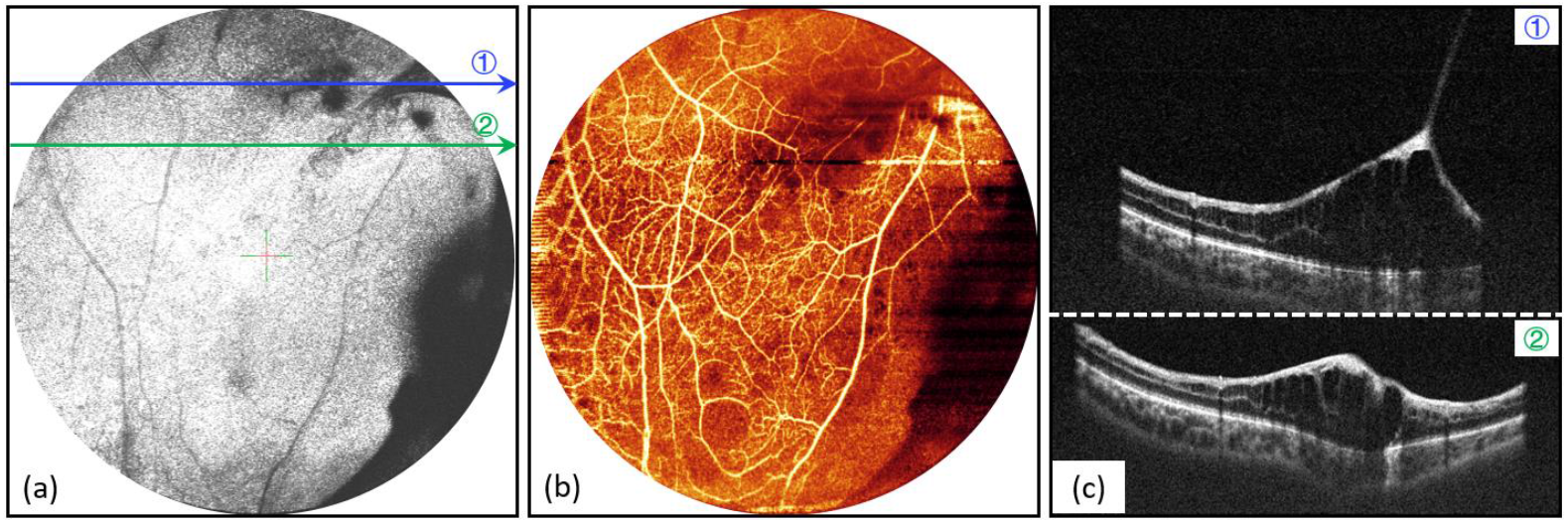
(a) *En face* OCT image from a patient with X-linked retinoschisis (XLRS) (b) Corresponding OCTA of the inner retina. (c) Two selected B-scans with the symptoms of retinal detachment.

Fig. 10 and Fig. 11 show the RetCam fundus, *en face* OCT image and OCTA in different areas from a 6-week-old baby with IP. IP is an X-linked dominant disease caused by mutations in the NEMO gene, which typically affects skin, hair, and retinovascular endothelial cells [38]. This can lead to vascular occlusion, neovascularization, optic nerve atrophy, and in later stages retinal detachment [39]. Similar to ROP, the preretinal neovascularization is typically found in the retinal periphery at the border of avascular retina [39,40]. The *en face* OCTA with smaller field of view shows more details in the area of interest (Fig. 11a and Fig. 11c). Fig. 11b and Fig. 11d show the neovascular area from the red box and green box of the RetCam fundus (Fig. 10a). Inner retina *en face* OCTA images were generated using maximum projection method, a customized purple color map was applied. Neovascular OCTA signals above the internal limiting membrane (ILM) layer and within the vitreous were also projected using a customized yellow color map and overlaid with the inner retina *en face* OCTA image. Vascular network and pathologies were visualized more clearly without pronounced motion artifacts.

**Fig. 10.**
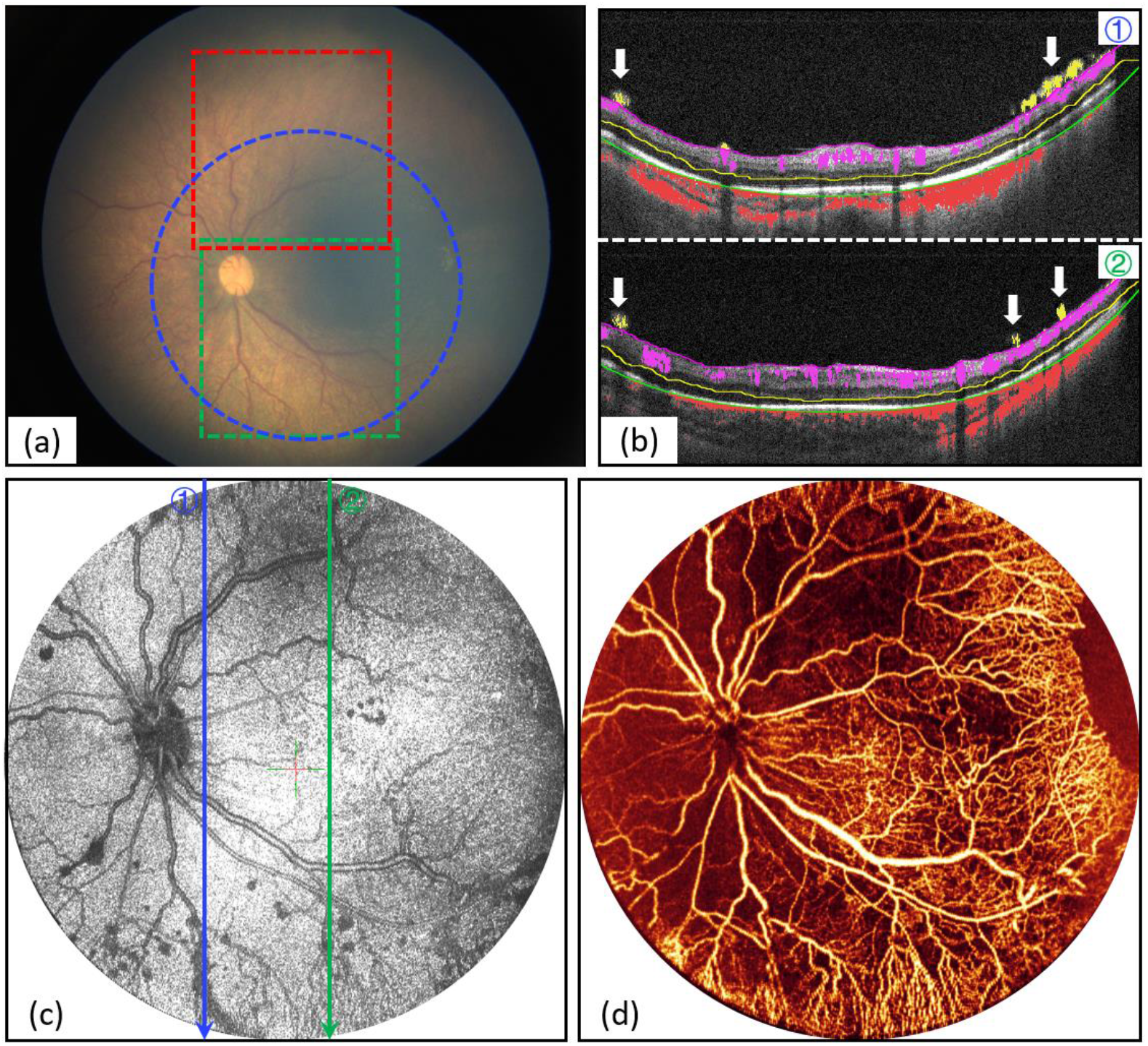
(a) RetCam fundus images of a 6-week-old baby presented with incontinentia pigmenti (IP). (b) Two selected B-scan reveals preretinal neovascularization (yellow vessels pointed by white arrows). (c) Handheld *en face* OCT image with 55-degree field of view in the blue circle of fundus photograph. (d) Corresponding OCTA of the inner retina.

**Fig. 11.**
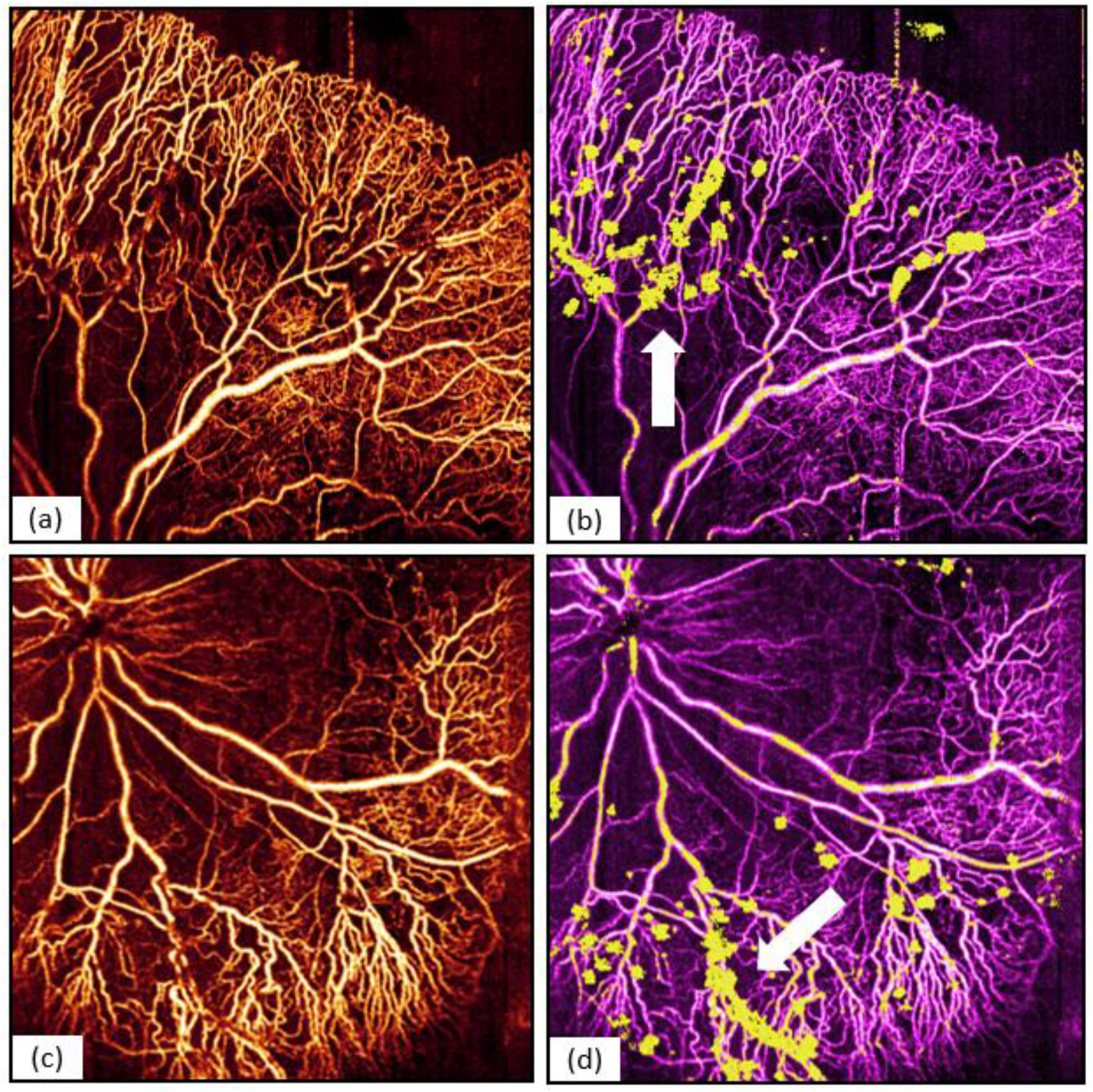
40-degree field of view *en face* OCTA of inner retina (a) and neovascular area (yellow vessels pointed by white arrow) (b) in the red box of fundus photograph (Fig. 10a). *En face* OCTA of the inner retina (c) and neovascular area (yellow vessels pointed by white arrow) (d) in the green box of fundus photograph (Fig. 10a).

## 4. Discussion

Aligning and imaging uncooperative, non-sedated, and non-fixating neonatal patients is difficult, and due to their medical comorbidities, it is critical to be able to obtain high quality images within 1-2 minutes. Thus, real-time visualization is a highly desirable function for a mission critical handheld imaging device and commercial handheld fundus cameras (RetCam) that are used routinely in ROP screening have implemented this function. Unlike conventional 2D imaging modalities, OCT requires optical path length matching between the reference and sample arms, which causes the alignment of OCT to be more challenging. Moreover, real-time OCT data processing and visualization is more computation intensive, and real time *en face* OCT requires the high speed OCT system to achieve usable refresh rate. Several excellent publications have demonstrated impressive progress on handheld OCT/OCTA with improved ergonomics and clinical utility [11,13,14,41–46]. The major effort in this study is to leverage the recent advance in laser light source, high performance computing, and optical design to improve a handheld OCT/OCTA system’s usability in pediatric retinal imaging. Compared to previous work, we were able to improve image quality while maintaining field of view and ergonomics, and reduce time required for imaging session.

With the combination of the two scanning modes and GPU accelerated OCT processing software, we believe we were able to balance the design trade-offs between imaging speed, field of view, and resolution. Warp scan mode has high *en face* imaging speed and is used for initial alignment and locating region of interest. OCTA mode has the large volumetric field of view and offers a panoramic view of the retinal layers and vasculature. Although our OCTA scan pattern does not satisfy the Nyquist sampling theorem, we were still able to acquire high quality OCTA images that provide sufficient angiographic information for disease diagnosis. We could increase the sampling density of the OCTA scans; however, it would result a longer imaging time and more severe motion artifacts. We found that with our current OCTA scanning pattern, motion artefacts caused by the operator hand tremor was minimal. The prevalent motion artefacts were introduced by the subjects’ eye motion. The 400-kHz VCSEL light source was instrumental in this study, as not only did it provide fast imaging speed, but also long depth range. Due to time constraints when imaging uncooperative infants, it is not always possible to adjust reference arm position so that the retina is placed close to DC. Therefore, having good imaging quality over the entire imaging depth is critical to improve imaging success rate.

The optical design of the OCT sample arm in this study significantly improved the optical performance in terms of controlling the system aberrations and minimizing the beam wander on the pupil plane. In the representative images shown in the results section, there were no apparent vignetting artefacts, and high quality OCTA signal was observed over the entire field of view, an evident of high-performance optical design as the OCTA signal is more sensitive to optical aberrations. Images with reduced field of view (40-degree, Fig. 11) showed exquisite details of the retinal capillaries and neovascularization indicating that our optical design provided sufficient transverse resolution down to capillaries level. The optimized interferometer increases the collection efficiency which is important in the high-speed OCT system. Our lightweight and ergonomic mechanical design allowed the operator to pivot the imaging probe to locate region of interest on the ocular fundus.

Despite the overall improved design and excellent OCTA image quality, there are some limitations of our handheld OCT imaging probe. The 55-degree field of view of our imaging system in this study is still not wide enough to cover the entire peripheral retina extending beyond zone 2 in one snapshot. An ultrawide field of view retinal imaging system with over 100-degree FOV is desired. OCTA with 100-degree FOV would require an even faster laser of over 1 MHz and a contact lens-based sample arm design, which cannot be assembled from off-the-shelf optical components. Our OCT imaging system cannot yet resolve all the capillaries, and further increasing the transverse resolution may require adaptive optics to compensate for the system and ocular aberrations especially at the peripheral retina [47,48]. And with high-speed swept-source OCT, a sensorless adaptive optics approach might be feasible in handheld imaging system [49–53]. It is also possible to average several OCTA *enface* images in order to improve the image quality and remove any residual motion artefacts [54]. The GPU OCT processing software and the on-probe display offer the operator real-time *en face* and cross-sectional views of the retina being imaged, but it does not display the *en face* OCTA images which could be more informative in disease diagnosis. Our GPU processing software is fast enough to calculate the OCTA B-scans, but real-time retinal layer segmentation is needed to generate meaningful *en face* OCTA images. We have demonstrated a deep learning based retina layer segmentation software that can segment 7 retina layers in an OCT B-scan in 3.5 ms [55], and we will improve and adapt this software for the handheld OCT in the future.

There are several compelling lessons learned from the adoption of OCT in adult retinal diseases that suggest OCT/OCTA may become essential not only for ROP but also for all pediatric retinal disease in the future. OCT is superior to ophthalmoscopic exams for the diagnosis of macular diseases and has changed the way diseases such as age-related macular degeneration (AMD) and diabetic retinopathy (DR), the two leading causes of vision loss in adults, are classified. OCT has demonstrated the ability to detect subclinical disease that not visible on ophthalmoscopy, enabling earlier treatment in both conditions. Quantitative OCT metrics have enabled objective disease monitoring. The development of OCT coincided with a therapeutic transition towards the use of anti-vascular endothelial growth factor (anti-VEGF) drugs, where OCT is indispensable for monitoring response to treatment and determining retreatment frequency. A similar transition is slowly occurring in ROP; however, the monitoring of treatment response and disease recurrence is an unsolved problem due to the lack of objective disease metrics. The widefield high speed handheld OCTA presented provides evidence that with real time visualization and high-speed, OCT/OCTA has the potential to be used as a screening tool for general pediatric retinal diseases.

## 5. Conclusion

We have demonstrated widefield (up to 55-degree) OCT and OCTA retinal imaging, with a 400-kHz handheld SS-OCT prototype that was custom designed for imaging infants. The novel optical design of the OCT sample arm significantly increases the scanning angle and minimizes the beam wander on the pupil plane without introducing excessive aberration. Our handheld imaging system overcame several obstacles in terms of imaging time constraints and system alignment to achieve good image quality. The OCT/OCTA images presented in this study clearly showed retinal structures and microvascular abnormalities. We believe that with the aid of our system, we can improve the capability of diagnosis and help make therapeutic decisions for pediatric retinal diseases.

